# Strigolactone defective mutants of *Arabidopsis* exhibit delayed sepal senescence

**DOI:** 10.1101/2020.01.07.897272

**Authors:** Xi Xu, Rubina Jibran, Yanting Wang, Lemeng Dong, Kristyna Flokova, Azadeh Esfandiari, Andrew McLachlan, Axel Heiser, Andrew Sutherland-Smith, Harro Bouwmeester, Paul Dijkwel, Donald Hunter

**Affiliations:** Massey University, School of Fundamental Sciences, Palmerston North, New Zealand; The New Zealand Institute for Plant and Food Research Limited, Palmerston North, New Zealand; Swammerdam Institute for Life Sciences, University of Amsterdam, Amsterdam, The Netherlands; AgResearch Limited, Hopkirk Research Institute, Palmerston North 4474, New Zealand.

**Keywords:** SLs, *Arabidopsis*, mutants, MAX1, AtD14, sepal senescence, darkness, sugar starvation

## Abstract

Flower sepals are critical for flower development and vary greatly in lifespan depending on their function postpollination. However, very little is known on what controls sepal longevity. Using a sepal senescence mutant screen, we directly connected strigolactones (SL) with sepal longevity. We identified two *Arabidopsis* mutants that harbour novel mutations in the SL biosynthetic gene *MORE AXILLARY GROWTH1* (*MAX1*) and receptor *DWARF14* (*AtD14*). The mutation in *AtD14* caused a substitution of the catalytic Ser-97 to Phe in the enzyme active site. The lesion in *MAX1* changed a highly conserved Gly-469 to Arg in the haem-iron ligand signature of the cytochrome P450 protein, which caused loss-of-function of MAX1. nCounter-based transcriptional analysis suggested an interaction between SL and sugar signalling in controlling dark-induced inflorescence senescence. The results uncover an important function for SL in regulating floral organ senescence in addition to its other diverse functions in plant development and stress response.

**One-sentence summary:** Two novel mutants in the strigolactone pathway demonstrate a role for the hormone in sepal senescence, and transcriptional analysis highlights interaction between strigolactones and sugar signalling.

## Introduction

Senescence typically occurs in mature cells of tissues after their growth phase has ceased to enable efficient recycling of nutrients to new growing sinks such as seeds (Thomas, 2013). At the whole plant level, senescence is considered critical for plant fitness, enabling plants to survive optimally in their given environments. The ability to senesce requires a change in competency of the tissue that happens during aging (Jing *et al*., 2005; Fracheboud *et al*., 2009). Nevertheless, imposition of stress can make tissue senesce early, possibly as an adaptive response to allow survival of the plant as a whole (Kanojia and Dijkwel, 2018). Prolonged darkness is a stress that results in carbon deprivation and early senescence of chlorophyllous tissues. This has been observed in attached leaves individually covered (Weaver and Amasino, 2001; Law *et al*., 2018), and in sepals of immature inflorescences of broccoli and *Arabidopsis* (Page *et al*., 2001; Trivellini *et al*., 2012). Research on the precocious senescence of these tissues has revealed that signalling key to their degreening is similar to that happening naturally in leaves *in planta* as they senesce in an age-dependent manner and in response to canopy shading. For example, the phytochrome interacting factors *PIF4* and *5* that have important roles in the shade response of canopy leaves (Sakuraba *et al*., 2014) were first identified to regulate the precocious degreening of harvested immature inflorescences of *Arabidopsis* held in the dark (Trivellini *et al*., 2012). Similarly, mutations in *EIN2* and *ORESARA1/ANAC092* that alter timing of sepal senescence of the dark-held inflorescences (Trivellini *et al*., 2012) were previously identified in *Arabidopsis* as components of the feedforward control of age-related leaf senescence (Kim *et al*., 2009).

Phytohormones have long been known to have key roles in senescence regulation. Ethylene, salicylic acid, abscisic acid, jasmonic acid and brassinosteroid promote the onset or progression of senescence; whereas cytokinin, gibberellic acid and auxin delay the process (Gan and Amasino, 1997; Lim *et al*., 2007). More recently, strigolactones (SLs), which are well-known for their function in regulating seed germination in parasitic plants (Toh *et al*., 2012; Wang and Bouwmeester, 2018) and plant shoot branching (Gomez-Roldan *et al*., 2008; Umehara *et al*., 2008), were reported to regulate natural- and dark-induced leaf senescence (Woo *et al*., 2001; Snowden *et al*., 2005; Yan *et al*., 2007; Hamiaux *et al*., 2012; Yamada *et al*., 2014; Ueda and Kusaba, 2015). Unsurprisingly, many of the regulators found to control senescence do so by regulating or being part of hormone biosynthetic or response pathways. For example, the MADS-box transcription factor *AGAMOUS-Like 42* is hypothesised to control senescence of the *Arabidopsis* flower *via* ethylene response factors (Chen *et al*., 2011). Abscisic acid influences dark-induced degreening of leaves (Noodén, 1988), age-related leaf senescence (Zhang *et al*., 2012) petal senescence (Panavas *et al*., 1998) and fruit ripening (Leng *et al*., 2014). Plant hormones also do not work alone, but rather in concert with others to control senescence progression. For example, ethylene, abscisic acid and jasmonates have been found to crosstalk with each other to control leaf senescence in *Arabidopsis* (Kim *et al*., 2011).

Very little is known on how sepal senescence is regulated in flowers with most research having focussed on petals. To identify key regulators of sepal senescence we systematically evaluated a population of mutant *Arabidopsis* plants derived from ethyl methanesulfonate (EMS) treated seeds. By using an *Arabidopsis* Inflorescence Degreening Assay (Hunter *et al*., 2018), we previously identified three independent mutations in chlorophyll *b* reductase that resulted in a delayed degreening phenotype (Jibran *et al*., 2015). Here we report on the characterization of a further two mutants with delayed senescence that highlight the role of SLs in controlling the lifespan of floral organs.

## Results

### Two EMS mutants exhibit delayed dark-induced degreening of excised immature inflorescences

To identify key genes that regulate dark-induced senescence, we used an *Arabidopsis* inflorescence degreening assay (Hunter *et al*., 2018). Using this assay, we screened ∼ 20,000 M2 plants grown from EMS–mutagenized seeds of *Arabidopsis* Landsberg *erecta* (L*er*-0) and identified 20 mutants, which had inflorescences that when excised and held in the dark showed delayed yellowing (termed *<u>d</u>elayed inflorescence <u>s</u>enescence*, *dis*) compared with wild-type (WT) (Jibran *et al*., 2015). Here, we report on two of these mutants, *dis9* and *dis15* (Fig. 1A). Their delayed degreening phenotypes were confirmed by chlorophyll (Chl) measurement at 3 days of dark incubation (Fig. 1B). Both mutants backcrossed to L*er*-0 WT produced 3:1 WT:mutant segregation ratios based on the delayed dark-induced degreening phenotype in their F2 populations (Supplementary Table S1). This indicated that a single locus was responsible for the *dis* phenotype in both mutants. The mutant plants were also dwarf and bushy compared with WT (Supplementary Fig. S1). These results suggested that the two *DIS* loci control degreening of excised immature inflorescence in the dark, flowering stem elongation and branching *in planta*.

**Fig. 1.**
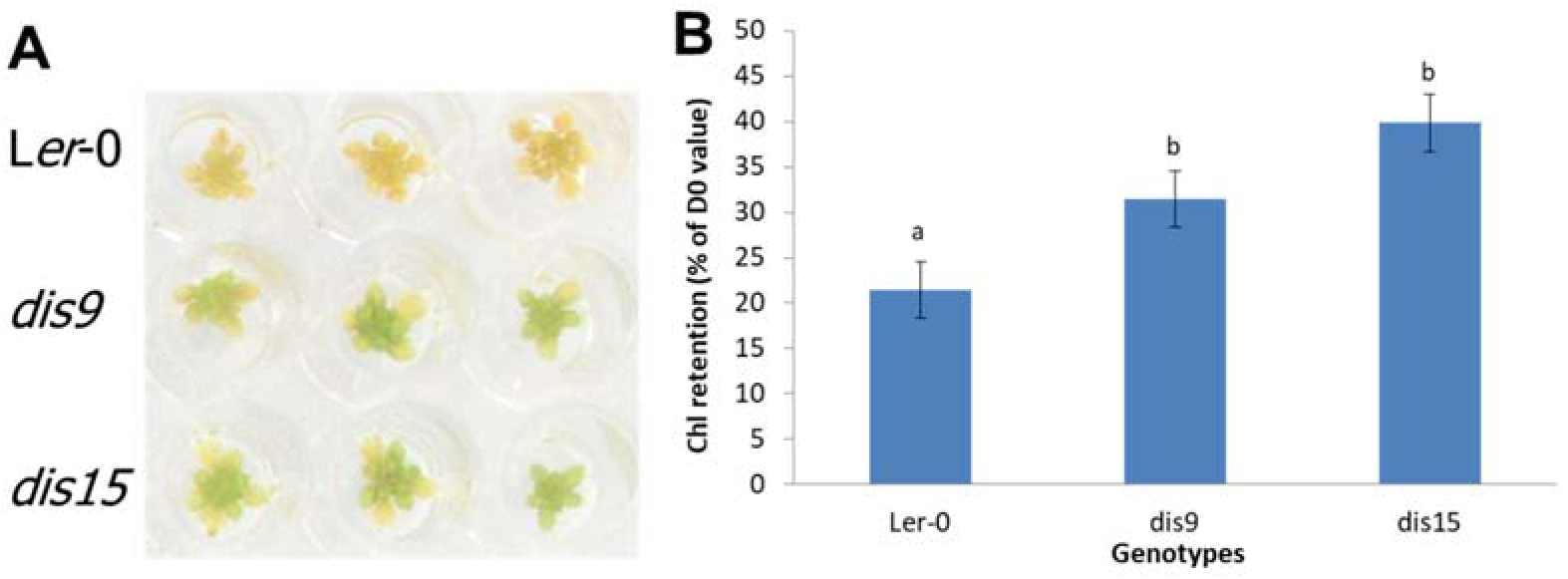
Two EMS mutants displaying delayed degreening of excised dark-incubated inflorescences. (A) Degreening of dark-incubated inflorescences. Inflorescences were excised and placed in water and incubated in the dark for 5 days. Three biological replicates are shown. (B) Total chlorophyll retention (% of day 0 value) at day 3 of dark incubation. Data are mean ± standard error (n=6). Letters represent significant differences between L*er*-0 and *dis* mutants in one-way ANOVA test (Fisher’s protected LSD test *P*<0.05).

### *dis9* and *dis15* also exhibit delayed sepal degreening *in planta*

Previous studies found that genes (e.g., *NYC1*, *NAP*, *ANAC092* and *ANAC046*) regulating dark-induced senescence also regulate natural senescence (Kim *et al*., 2018b). Therefore, we hypothesized that the DIS loci in both mutants also controls sepal degreening during plant development. To test this, we grew plants for 4 – 5 weeks in long day conditions (16 h/8 h light/dark cycles) to allow the floral organs to develop and observed the colour of the sepals when they were starting to abscise. In all four independent experiments, the sepals of the mutants were always green when they abscised. In comparable WT plants, they were yellow (Fig. 2A & B; Supplementary Fig. S2A). The delayed sepal yellowing also occurred in the detached inflorescences that were incubated in long day conditions (Fig. 2C; Supplementary Fig. S2B). This indicates that in addition to affecting the timing of dark-induced degreening, the *DIS* loci also controls sepal degreening *in planta* and in detached inflorescences held in long days.

**Fig. 2.**
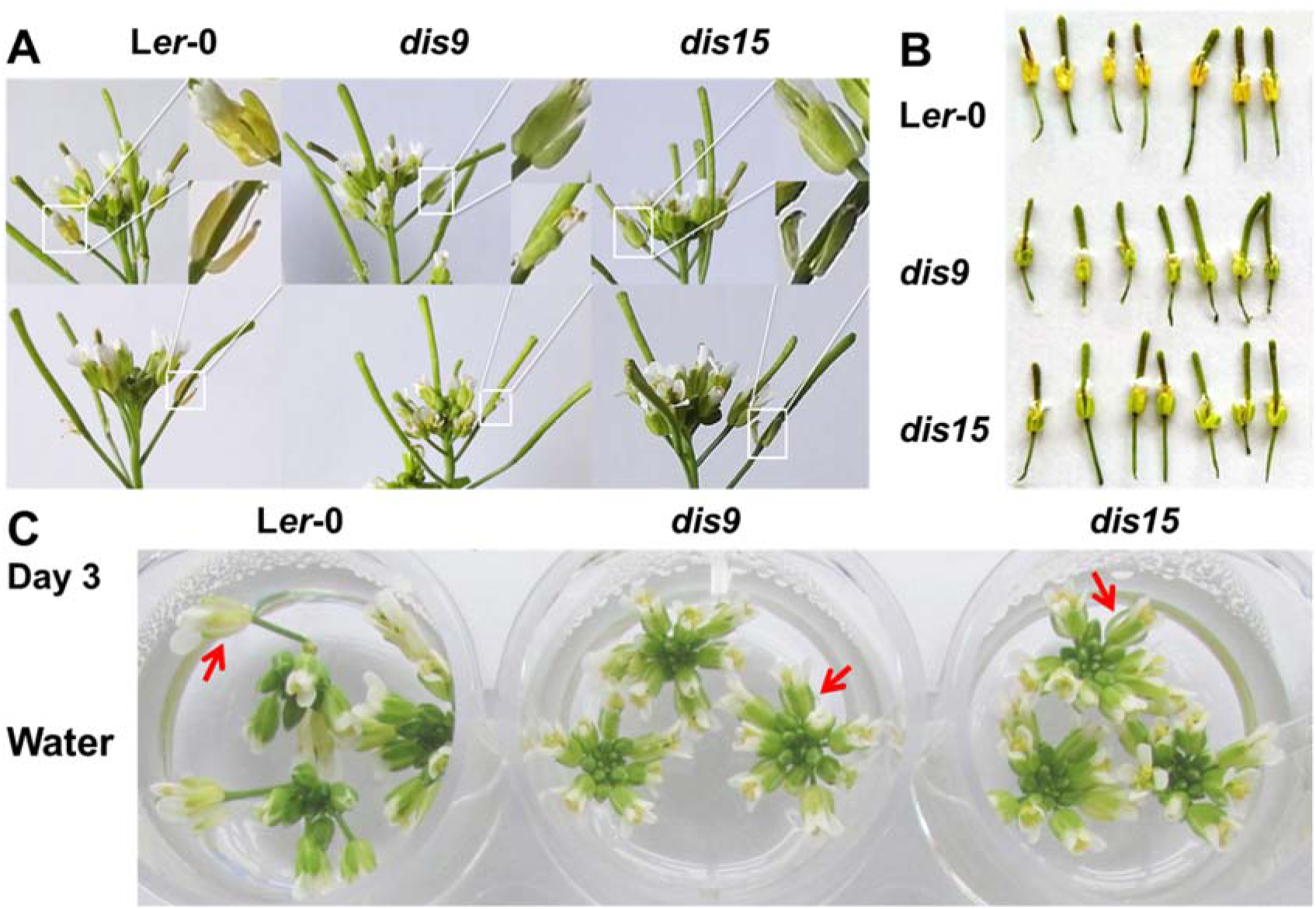
Delayed sepal degreening of *dis9* and *dis15 in planta* and in detached immature inflorescences held in long day conditions (16 h/8 h light/dark cycle) (A) Inflorescences attached to plants. Inflorescences from the primary bolts of 4.5-week-old wild type and *dis* plants were photographed. Two biological replicates with representative abscising sepals (circled in white and magnified) are shown. (B) Degreening of sepals *in planta*. Flowers of 5-week-old plants were harvested when their sepals were just starting to abscise. Each flower of the seven biological replicates is from an independent plant. (C) Sepal degreening of excised inflorescences. Inflorescences were harvested from the primary bolts of 4.5-week-old plants that had their first flower opened on the same day. The inflorescences with removed opened buds were placed in water and incubated in 16h/8h light/dark cycle for 3 days. Three biological replicates are shown. Representative sepals are indicated by red arrows.

### Transcript abundance of senescence marker genes are suppressed in *dis9* and *dis15* during dark incubation

Dark-induced senescence is associated with increased transcript accumulation of senescence associated genes (SAGs) (Buchanan-Wollaston *et al*., 2005; Trivellini *et al*., 2012) and some such as *SAG12* and *ANAC092* have often been used as indicators of senescence progression (Grbic, 2003; Kim *et al*., 2009; Balazadeh *et al*., 2010; Trivellini *et al*., 2012). Therefore, we compared the transcript abundance of these marker genes in WT and the two mutants during continuous dark treatment. The transcript abundance of the genes increased substantially upon dark treatment in both L*er*-0 WT and mutants, but the increase in the mutants was significantly less than that of WT at 72 h (Fig. 3). This suggests that senescence progression in the mutants was suppressed compared with that in WT.

**Fig. 3.**
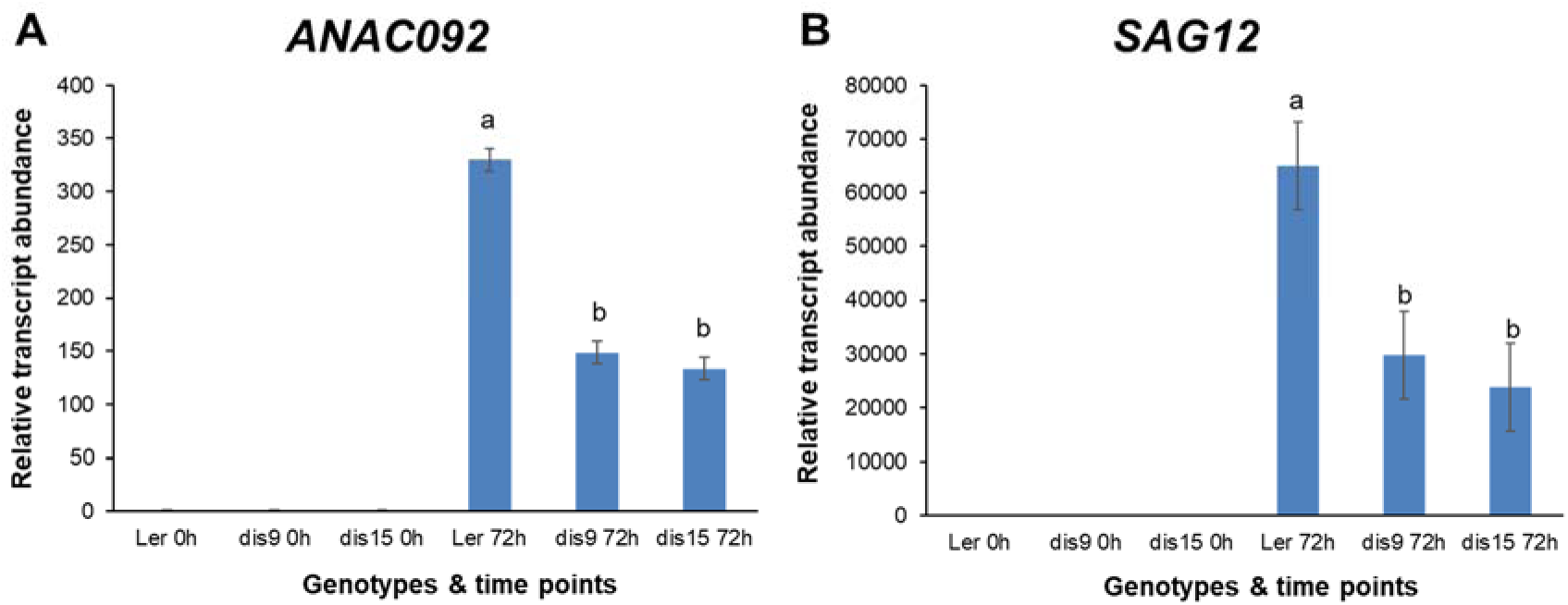
Transcript abundance based on qPCR analysis of *ANAC092* and *SAG12* in L*er*-0 and *dis* mutants at 0 h and 72 h. (A) for *ANAC092*. (B) for *SAG12*. In both (A) and (B), transcript abundance was normalised to *PP2AA3* and expressed relative to L*er*-0 0 h. Data are mean ± standard error (n=3). Letters represent significant differences between L*er*-0 and *dis* mutants at 72h dark incubation for each gene comparison in one-way ANOVA (Fisher’s protected LSD test *P*<0.05). Means with same letter denote not significantly different between *dis9* and *dis15*.

### *dis9* and *dis15* have genetic lesions in *AtD14* and *MAX1* respectively

To identify the genetic lesions in *dis9* and *dis15*, the mutants were outcrossed to Col-0 WT. F2 progeny from both crosses segregated based on the delayed dark-induced degreening phenotype at 3:1 WT:mutant suggesting the mutations were monogenic recessive (Supplementary Table S1). The *dis9* mutation was mapped by HRM analysis to the top arm of Chromosome 3 (Chr 3) between 0.6-1.5 Mbp (Supplementary Fig. S3A), and *dis15* was mapped to the bottom arm of Chr 2 between 9.6-12.4 Mbp (Supplementary Fig. S3B). Whole genome sequencing (WGS) analysis of DNA from 50 pooled *dis9* positive plants from a segregating population identified a C to T transition at position 290 downstream of the translation start site (TSS) of the *Arabidopsis DWARF14* (*AtD14* gene, AT3G03990). This mutation substituted a Phe for Ser at position 97 (S97F) in the AtD14 α/β-fold hydrolase protein that functions as a SL receptor (Arite *et al*., 2009; Yao *et al*., 2016). WGS data from *dis15* positive M4 plants revealed a G to A transition at position 1405 downstream of the TSS of the coding sequence of the *MORE AXILLARY GROWTH1* (*MAX1* gene, AT2G26170). This mutation resulted in a substitution of Gly to Arg at position 469 (G469R) in the MAX1 cytochrome P450 monooxygenase (Booker *et al*., 2005) that is involved in SL biosynthesis converting carlactone (CL) to carlactonoic acid (CLA) (Abe *et al*., 2014).

### Treatment with rac-GR24 rescues delayed sepal degreening of *dis15* but not *dis9*

To test whether the sepal degreening phenotype of *dis15* was caused by lack of SL synthesis, we investigated whether exogenously applied SL could chemically complement the *dis15* sepal degreening phenotype. To do this, inflorescences of *dis15, dis9* and L*er*-0 WT were treated with different concentrations (1, 5, 25, 50 µM) of the SL analog rac-GR24. rac-GR24 can also induce karrikin (KAR) signalling, but this is likely not relevant for the dark-induced degreening phenotype as defects in KARRIKIN INSENSITIVE 2 (KAI2), a KAR-specific receptor, do not delay dark-induced leaf senescence (Ueda and Kusaba, 2015). rac-GR24 at 5 µM was sufficient for rescuing the delayed sepal degreening phenotype in *dis15* but not *dis9* at day 5 of dark incubation (Fig. 4A; Supplementary Fig. S4). Under long day (16-h light) conditions, the hormone analogue also hastened sepal yellowing of *dis15* but not *dis9* excised inflorescences (Fig. 4B; Supplementary Fig. S5). These results are consistent with the delayed sepal degreening phenotypes of *dis9* and *dis15* being caused by defects in SL signalling and biosynthesis, respectively.

**Fig. 4.**
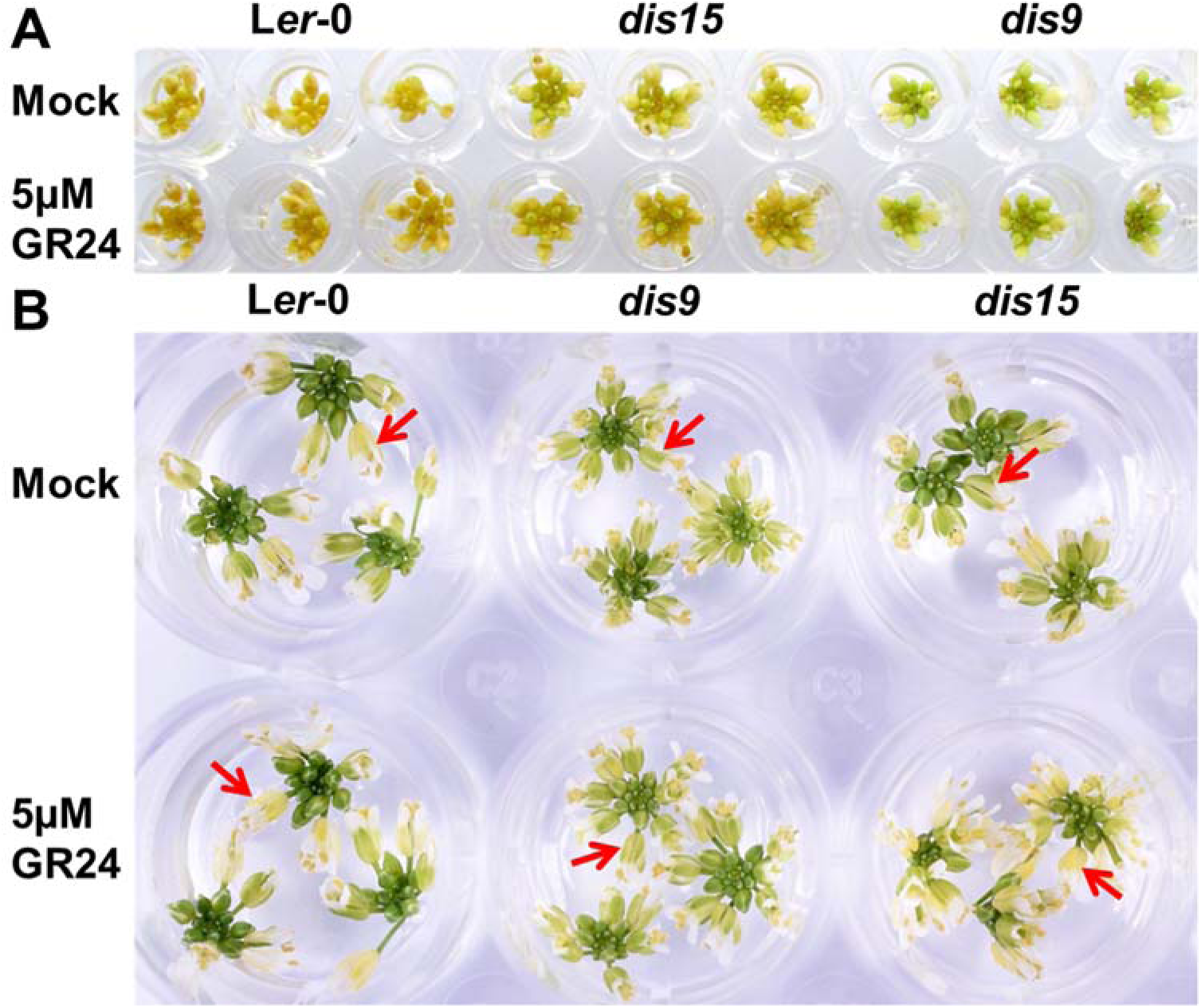
Sepal degreening of excised inflorescences treated with rac-GR24. (A) Dark incubated inflorescences. Inflorescences were harvested from the primary bolts of 4.5-week-old plants that had their first flower opened on the same day. The inflorescences with removed opened buds were treated with 1% DMSO (mock) and 5 µM rac-GR24 and incubated in the dark for 5 days. Three biological replicates for each genotype and treatment are shown. (B) Long day (16 h light) treated inflorescences. Inflorescences were harvested from the primary bolts of 4.5-week-old plants that had their first flower opened on the same day. The inflorescences with removed opened buds were treated with 1% DMSO (mock) and 5 µM rac-GR24 and incubated in 16h/8h light/dark cycle for 3 days. Red arrows indicate representative sepals. Three biological replicates are shown.

### Degreening phenotypes of *dis9* and *dis15* are complemented with wild type *AtD14* and *MAX1* genomic regions respectively

We confirmed that the mutations in the two SL-associated genes were responsible for the dis phenotypes by complementation. The *dis* phenotype of *dis9* was complemented with a WT *AtD14* construct (Supplementary Fig. S6) that previously rescued the increased branching phenotype of the *Arabidopsis d14-2*/*seto5* mutant (Chevalier *et al*., 2014). The *dis15* plants were complemented with a WT *MAX1* construct consisting of the 1293 bp genomic region upstream of the start codon, the 2420 bp *MAX1* genomic sequence, and a 185 bp region downstream of the stop codon (Supplementary Fig. S7). In summary, the delayed sepal degreening of *dis9* and *dis15* was caused by non-synonymous nucleotide changes in the coding sequences of *AtD14* and *MAX1,* respectively. These two mutants were therefore renamed as *d14-6*/*dis9* and *max1-5*/*dis15* respectively.

### Highly conserved amino acids are substituted in *d14-6*/*dis9* (S97F) and *max1-5*/*dis15* (G469R)

The striking delayed senescence and bushy phenotypes of *d14-6*/*dis9* and *max1-5*/*dis15* suggested that residues Ser-97 in D14 and Gly-469 in MAX1 were essential for proper functioning of their respective proteins. The D14 S97F substitution in *dis9* occurred at the Ser-His-Asp catalytic triad that is located at the hydrophobic substrate-binding pocket of the D14 hydrolase. Ser-97 in D14 of *Arabidopsis* is highly conserved in the orthologs from other species (Challis *et al*., 2013), including the characterised functional orthologues in petunia *Ph*DAD2 (Hamiaux *et al*., 2012), rice *Os*D14 (Arite *et al*., 2009; Gao *et al*., 2009; Liu *et al*., 2009), poplar *Pt*D14a (Zheng *et al*., 2016), pea RMS3 (de Saint Germain *et al*., 2016) and the paralog *At*KAI2 (Waters *et al*., 2012) (Fig. 5A). Changing the Ser-97 residue to alanine (a non-nucleophilic residue) was previously reported to cause loss of D14 hydrolase activity (Abe *et al*., 2014) and prevent formation of a covalently linked intermediate molecule (CLIM) in the active site of the protein (Yao *et al*., 2016) (Fig. 5B). The SL-defective phenotype we observed strongly suggests that substitution of Ser-97 to Phe (also a non-nucleophilic amino acid) caused loss of receptor activity *in planta*.

**Fig. 5.**
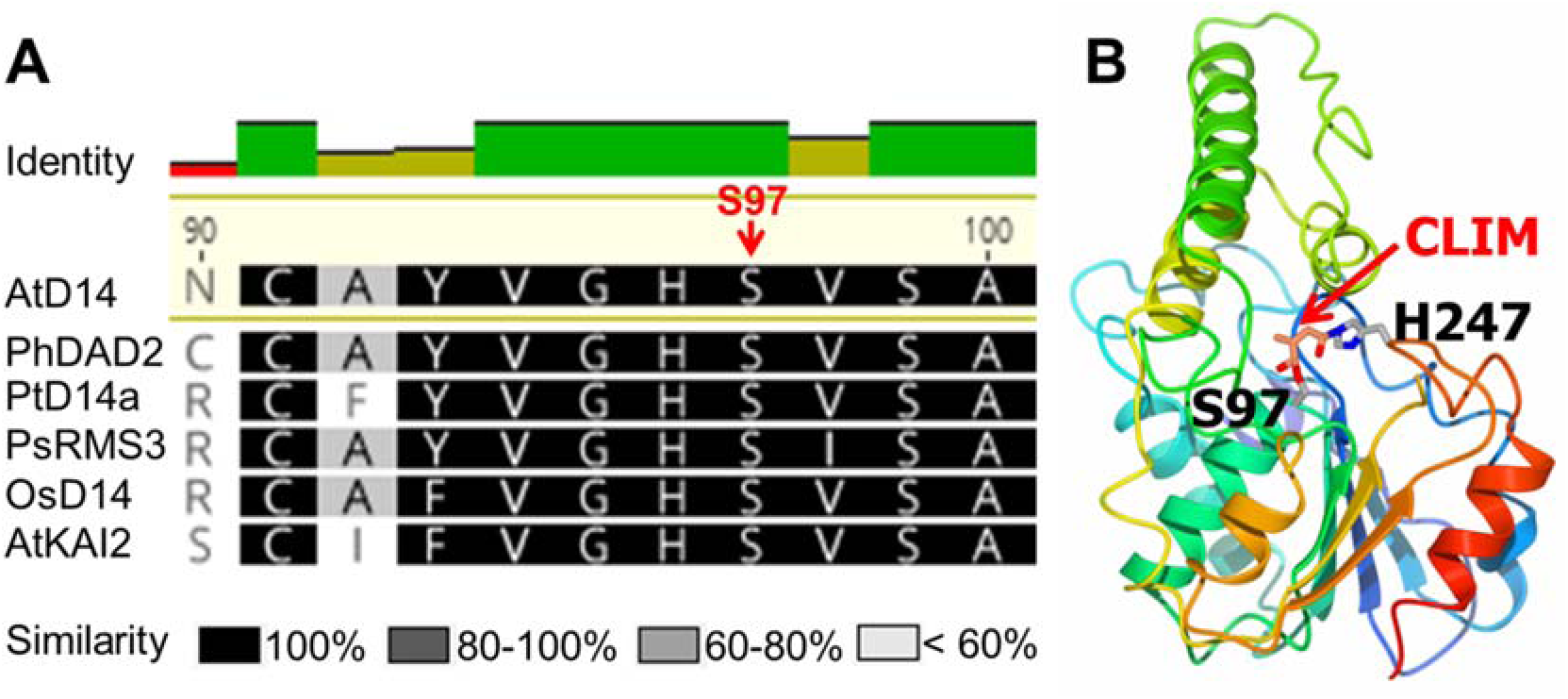
Location of the serine to phenylalanine substitution in the D14 protein. (A) Sequence alignment of *Arabidopsis* D14 with characterised homologs. Position of the mutation at Ser-97 (S97) in AtD14 is indicated in red. Amino acid positions are based on Col-0 sequence from TAIR. Aligned sequences were sorted by the differences to AtD14 reference sequence. Intensity of shading represents the percentage similarity of each residue among D14 homologs. At, *Arabidopsis*; Os, rice; Ph, petunia; Pt, poplar; Ps, pea. (B) Structure of AtD14 from the SL-induced AtD14-D3-ASK1-complex (PDB: 5HZG). The CLIM is shown as orange and red sticks. The catalytic triad residues Ser-97 and His-247 are shown in atomic colouring as grey/blue/red sticks. AtD14 is shown in cartoon representation coloured in a rainbow scheme (N to C terminus from blue to red).

The MAX1 G469R substitution in *max1-5*/*dis15* occurs in the last residue of the cysteine haem-iron ligand signature [FW]-[SGNH]-x-[GD]-{F}-[RKHPT]-{P}-C-[LIVMFAP]-[GAD], which is highly conserved in the cytochrome P450 superfamily (Prosite: https://prosite.expasy.org/PDOC00081). This G469 residue is invariant in all MAX1 functional orthologues studied thus far (Fig. 6A) (Yoneyama *et al*., 2018). However, as the ligand signature [GAD] indicates, glycine (G) can be replaced by alanine (A) or aspartate (D). This occurs at very low frequency in the wider cytochrome P450 protein family with G replaced by A in 3.4% or by D in 0.18% of the 1087 predicted cytochrome P450 proteins that have the cysteine haem-iron ligand pattern (according to the Prosite database) (Supplementary Fig. S8). We modelled the 3D structure of AtMAX1 on the most closely related cytochrome P450 (human cytochrome P450 CYP3A4) (Yano *et al*., 2004) with a protein crystal structure available (sequence identity of 28% and E value of 7e-51). The model showed that G469 is in the haem pocket packed against the haem group and is close to the haem-iron ligand cysteine (C467) (Fig. 6B). The G469R substitution may cause loss-of-function of *max1-5*/*dis15* by disrupting the steric structure of this pocket because there is not enough space in this pocket to accommodate Arg, which has one of the largest side-chains, compared with Gly, which has the smallest one.

**Fig. 6.**
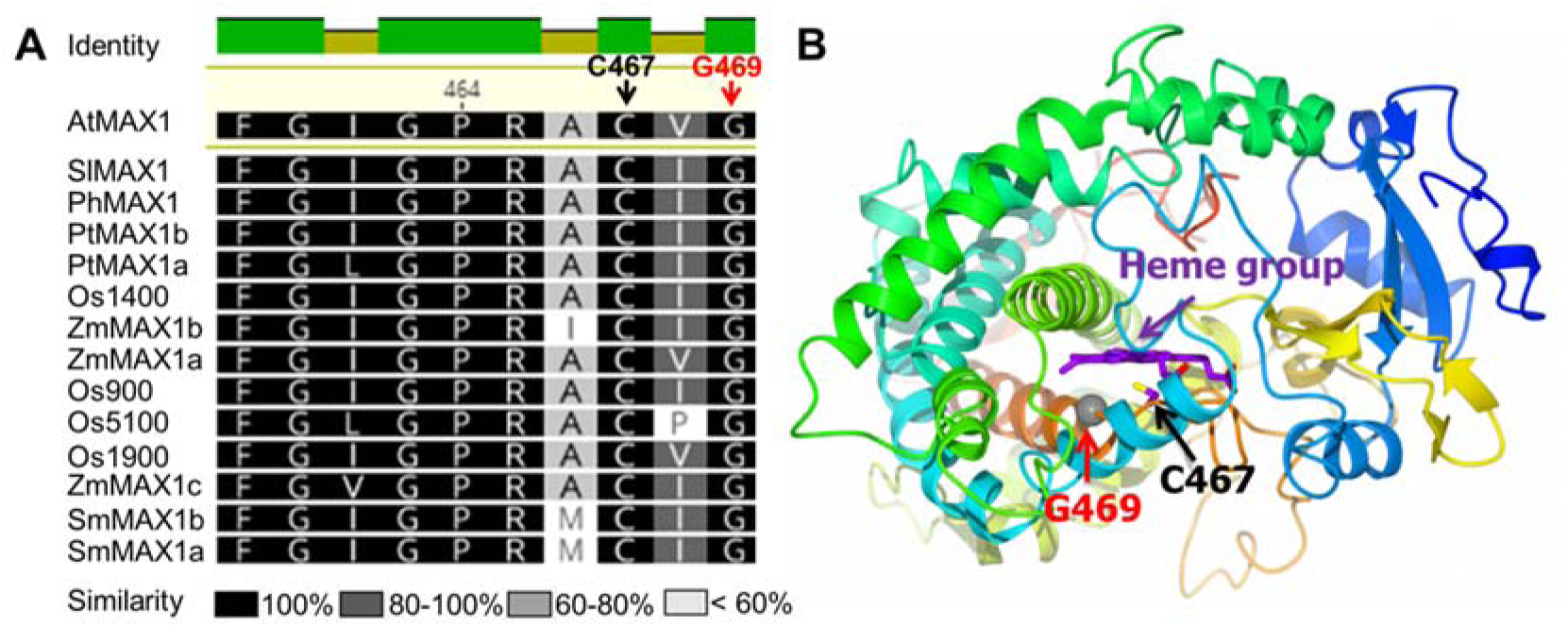
Location of the glycine to arginine mutation in the MAX1 protein. (A) Sequence alignment of *Arabidopsis* MAX1 with its functional orthologs. Position of mutation at Gly-469 (G469) and of the heme-iron ligand at Cys-467 (C467) in *Arabidopsis* MAX1 is indicated in red and black, respectively. Amino acid positions are based on Col-0 sequence from TAIR. The sequences of the cysteine haem-iron ligand signature are shown. Aligned sequences were sorted by the differences to AtMAX1 reference sequence. Intensity of shading represents the percentage similarity of each residue among MAX1 orthologs. At, *Arabidopsis*; Sl, tomato; Sm, *Selaginella*; Os, rice; Ph, petunia; Pt, poplar; Zm, maize. (B) MAX1 modelled on the structure of the closely related human cytochrome P450 CYP3A4 (PDB: 1TQN). The haem group is presented as purple sticks (carbons mostly), which is indicated by a purple arrow. The side chain of Cys-467 (in black; haem ligand) is represented by sticks (carbon is purple and sulphur is yellow). The position of Gly-469 is indicated by the grey sphere and is highlighted by a red arrow. The structure is presented in cartoon representation coloured in a rainbow scheme (N to C terminus from blue to red).

In summary, the S97F substitution in *d14-6*/*dis9* occurs at the catalytic centre of the D14 hydrolase, which likely causes loss-of-activity of D14 because the nucleophilic Ser was replaced by the non-reactive Phe. In *max1-5*/*dis15*, the G469R mutation in MAX1 occurs in the highly conserved haem-iron ligand signature of cytochrome P450 enzymes, which likely leads to a non-functional MAX1 protein by disrupting the steric structure of the haem-iron binding pocket.

### G469R substitution in *max1-5*/*dis15* disrupts enzyme activity of MAX1

To confirm the loss of activity of *max1-5*/*dis15* (MAX1-G469R) suggested by *in silico* prediction, we used transient expression in *N. benthamiana* developed to study the function of SL biosynthetic enzymes (Zhang *et al*., 2014). *Arabidopsis MAX1*-WT, *MAX1*-G469R and *MAX1*-G469A were transiently expressed with the upstream enzyme-encoding genes of the CL biosynthetic pathway (*AtD27*, *AtMAX3*, and *AtMAX4*) in *N. benthamiana* and the substrate (CL) and product (CLA) of MAX1 were measured. Transient expression of *AtD27*, *AtMAX3*, and *AtMAX4* indeed resulted in the production of CL and not CLA (Fig. 7A). When co-expressed with *MAX1*-WT (either L*er*-0 or Col-0 version) or with *MAX1*-G469A, CL was significantly reduced and some CLA was detected. However, when co-expressed with *MAX1*-G469R, the amount of CL did not decrease although CLA was produced in similar amounts as that produced by *MAX1*-WT (Fig. 7A). The lack of a decrease in CL suggested that MAX1-G469R had reduced enzymatic activity. We considered that the absence of a difference in CLA production in the different treatments was likely caused by conjugation (for example glycosylation) of CLA by endogenous *N. benthamiana* enzymes, as we had observed this several times before in *N. benthamiana*, e.g., in the transient production of geranic acid that was glycosylated with one or two hexoses (Dong *et al*., 2013). If CLA conjugation occurs efficiently, the amounts of free CLA would remain low and not reflect the rate of conversion of CL to CLA. Thus, we investigated whether *N. benthamiana* leaves expressing *MAX1*-WT accumulated CLA conjugates using LC-QTOF-MS analysis. Indeed, *N. benthamiana* leaves expressing the CL pathway genes together with *MAX1*-WT accumulated CLA-dihexose and CLA-hexose conjugates (Figure 7B, Supplementary Fig. S9). When *MAX1*-G469A was substituted for *MAX1*-WT, conjugate formation was not significantly different from that in the pathway with *MAX1*-WT (Figure 7B, Supplementary Fig.S9). However, when *MAX1*-G469R was substituted for *MAX1*-WT conjugate production decreased 36- and 15-fold, respectively. To confirm that the G469R mutation was affecting enzyme activity rather than exerting its effect through transcriptional changes, mRNA abundance of *MAX1*- WT and *MAX1*-G469R was analysed. There was no difference in expression between *MAX1*-WT and *MAX1*-G469R when they were expressed in *N. benthamiana* (Supplementary Fig. S10). Thus, we conclude that the lack of CL conversion and reduced CLA conjugate production in *N. benthamiana* upon co-infiltration of the CL pathway with *MAX1*-G469R was caused by loss-of-activity of the MAX1-G469R enzyme.

**Fig. 7.**
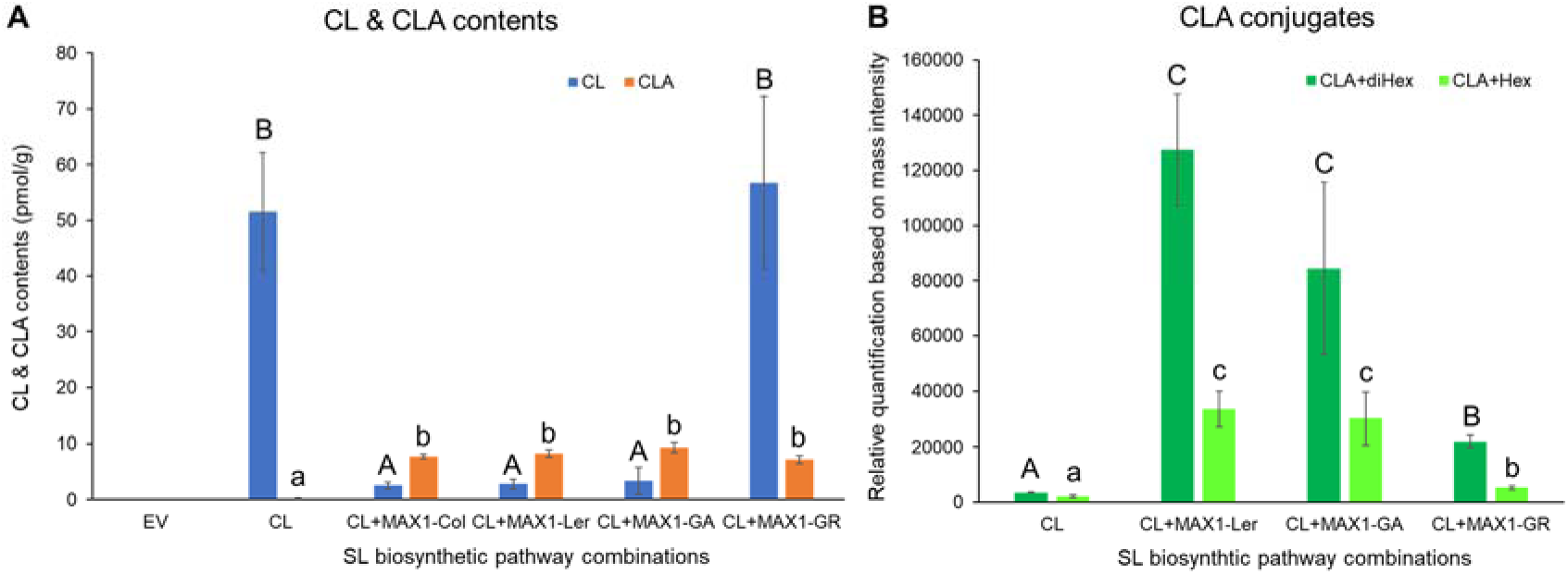
Analysis of CL, CLA and CLA conjugates in *N. benthamiana* leaves infiltrated with strigolactone biosynthetic gene constructs. (A) CL and CLA content in *N. benthamiana* transiently expressing *MAX1* (Col/L*er*-WT or with nucleotide changes resulting in G469R or G469A substitutions) plus CL pathway genes. Data are mean ± standard error (n=6). (B) Identification of CLA conjugates in *N. benthamiana* transiently expressing *MAX1*- L*er*/G469R/G469A plus CL pathway genes. Data are mean ± standard error (n=3). EV, empty vector (control); CL, carlactone pathway (expressing *AtD27*, *AtMAX3*, and *AtMAX4*); *MAX1*-Col/L*er*, CL pathway plus *MAX1* based on Col-0 or L*er*-0 sequence; *MAX1*-GR/GA, CL pathway plus *MAX1* with G469R or G469A substitutions based on L*er*-0 sequence. Letters represent significant differences among different gene combinations for the infiltration for each compound comparison in one-way ANOVA. Upper and lower cases were used to distinguish the difference for each compound in (A) and (B). Means for the same compound with the same letter are not significantly different (5% least significant difference comparisons made on log-transformed data).

### MAX1 remains functional when Gly-469 is replaced by Ala

As mentioned above, Gly-469 in MAX1 is substituted by Ala in 3.4% of the 1087 predicted cytochrome P450 proteins, but is invariant in MAX1 orthologues. Nevertheless, above we show that Ala can functionally replace Gly-469 (Fig. 7, Supplementary Fig. S9). The ability of Ala to functionally replace Gly-469 was also supported by *MAX1*-G469A complementing the *dis* phenotype of the *max1-5*/*dis15* mutant. (Supplementary Fig. S7). Together, the results support the prediction from *in silico* analysis that the G469R mutation makes MAX1 non-functional by physically disrupting the steric structure of the haem-iron binding pocket.

### SL biosynthetic and response genes are up-regulated by 24 h of dark incubation in inflorescences

Our data above showed that SLs participate in controlling inflorescence sepal senescence. We then asked at what stage of senescence SLs were involved. It has been suggested that the SL biosynthetic pathway is induced by senescence signalling (Ueda and Kusaba, 2015). If so, SL pathway genes would be expected to be induced later than senescence marker genes. To test this, we compared the timing of transcriptional changes in selected senescence-marker and SL-pathway genes in excised WT inflorescences every 24 h over a period of 3 days of dark treatment using nCounter Technology (Fig. 8). Transcript abundance of early stage senescence markers, i.e. Chl degradation gene *SGR1* (Park *et al*., 2007) and central regulator of senescence *ANAC092* (Balazadeh *et al*., 2010) were significantly increased at 24 h (Fig. 8A), suggesting senescence in the inflorescences had already initiated by this time. Increased transcript abundance of the late stage senescence-specific marker gene *SAG12* (Grbic, 2003) at 48 h indicated that at 2 days of dark incubation senescence was well advanced.

**Fig. 8.**
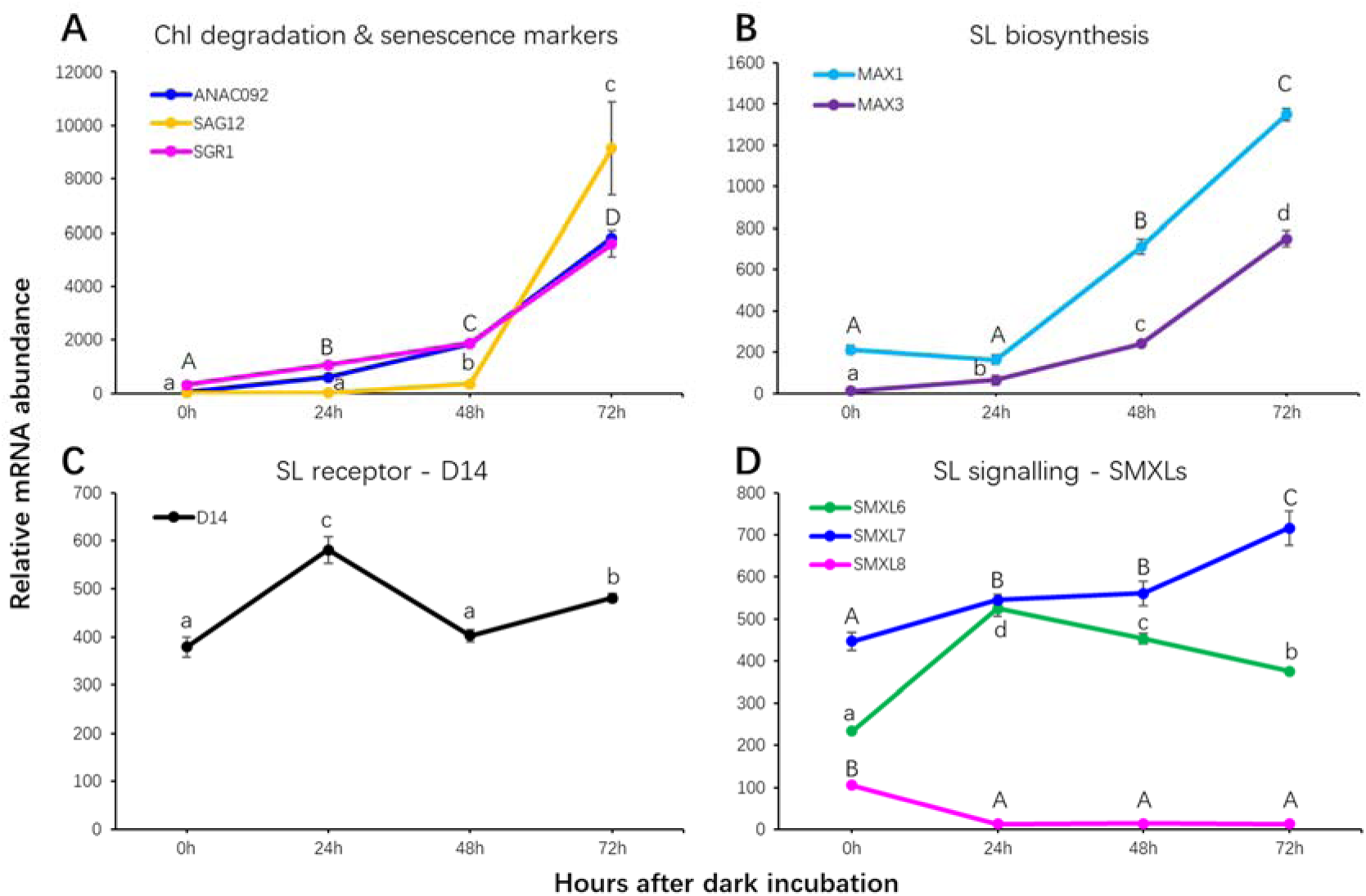
Dark-induced transcript abundance changes of strigolactone pathway and senescence-related genes. (A) Chlorophyll (Chl) degradation and senescence marker genes; (B) Strigolactone (SL) biosynthetic genes; (C) SL receptor; (D) SL signalling genes. Transcript abundance was quantified using nCounter technology on RNA isolated from detached WT inflorescences (n = 3 samples, > 4 inflorescences from independent plants per sample) that were incubated in the dark for 0 h, 24 h, 48 h and 72 h. Transcript abundance was normalised to geometric mean of *PP2AA3*, *ACT2* and *MON1*. Data are mean ± standard error. Letters represent significant differences among four time points for each gene comparison in one-way ANOVA (Fisher’s protected LSD test *P*<0.05). Upper and lower case letters were used to distinguish the comparisons for each gene. Upper case letters in (A) represented comparisons for both *ANAC092* and *SGR1*.

We used the transcript abundance changes of three SL biosynthetic genes *MAX1*, *MAX3* and *MAX4* to determine when SL biosynthesis had initiated in the dark-held inflorescences. *MAX1* transcript abundance did not significantly change during the first 24 h of dark treatment, but then significantly and substantially increased to be highest at 72 h (Fig. 8B). *MAX3* transcript abundance was slightly increased at 24 h, suggesting that SL production in the tissue was just starting. From 24 h onwards, *MAX3* transcript abundance increased in concert with both early senescence markers *SGR1* and *ANAC092*. qRT-PCR analysis of *MAX4* revealed a pattern of transcript accumulation that was very similar to that of *MAX3* suggesting co-regulation and beginning of SL synthesis by 24 h (Supplementary Fig. S11A).

The timing of SL response was determined by measuring changes in transcript abundance of the SL signalling genes *AtD14*, *SMXL6*, *SMXL7* and *SMXL8*. Transcript abundance of the first three genes increased significantly at 24 h of dark treatment (Fig. 8C, D), whereas *SMXL8* transcript abundance decreased to be undetectable at 24 h (Fig. 8D). The nCounter results for *MAX3*, *SMXL6/8*, *ANAC092* and *SAG12* were confirmed by qRT-PCR analysis (Supplementary Fig. S11B, C).

Overall the results from the transcript profiling of the inflorescence suggest that by 24 h of dark incubation SL biosynthesis has been initiated, SL signalling is occurring, and senescence has started. Thus, earlier time points were investigated to determine the order of pathway activation.

### SL signalling but not biosynthetic genes respond rapidly to the light-dark transition

At 24 h of darkness, the inflorescence tissue had been exposed to 8 h of regular and 16 h of extended night. One of the most notable outcomes of keeping tissue in extended darkness for this length of time is acute carbon starvation caused by exhaustion of starch reserves (Usadel *et al*., 2008), which causes precocious senescence. This agrees with our previous work that found that carbon-depletion resulting from a 24 h dark treatment is a key senescence stimulus for detached immature inflorescences (Trivellini *et al*., 2012). We therefore considered the possibility that SL biosynthesis and response early in the extended night are associated with acute carbon deprivation-based signalling. To test this, we compared the timing of expression of transcriptional markers of tissue carbon status (SnRK1-related genes *AKINβ1* and *bZIP63*) (Blasing *et al*., 2005; Usadel *et al*., 2008; Li *et al*., 2009; Mair *et al*., 2015) with that of SL-associated genes both in the regular and early extended night.

In WT, during the first 6 h into the regular night, transcript abundance of the *AKINβ1* and *bZIP63* increased in the detached inflorescences held in the dark (Supplemental FigS12A). This was consistent with them being markers of reduced (but not yet acutely limited) carbon availability. Key senescence-regulatory genes *ANAC092* (Kim *et al*., 2009) and *AtNAP* (Guo and Gan, 2006) that are also sugar responsive were upregulated at 3 h (Supplemental FigS12B). Since these two genes are also diurnally regulated (Kim *et al*., 2018a; Song *et al*., 2018), their transcriptional increase was more likely in response to changes in soluble sugar (Blasing *et al*., 2005; Usadel *et al*., 2008) and circadian clock rather than senescence. Because of this, we did not expect SL biosynthesis and signalling genes to respond during this early night time frame. As expected, during the first 6 h into the regular night there was no increase in transcript abundance of *MAX1*, while *MAX3* transcripts were not detected (Fig. 9A). Transcript abundance of *SMXL7* also did not change during the regular night (Fig. 9B). However, at 3 h into the dark period *SMXL6* and *SMXL8* were up- and down- regulated, respectively (Fig. 9B). In the *max1-5*/*dis15* mutant, the two *MAX* and three *SMXL* genes exhibited similar expression patterns to that of WT, although their abundance was lower (Fig. 9A, B). The reduced expression of the three *SMXL*s was reversed when the mutant was treated with rac-GR24 for 3 h (Fig. 9B; Supplementary Table S2). Intriguingly, *AtNAP* was significantly upregulated by rac-GR24 at 3 h of treatment, suggesting it is also a SL-inducible gene. Thus, based on transcription, the WT inflorescences experienced a reduction in sugar content during the normal night, but this did not trigger initiation of SL biosynthesis. In the *max1-5*/*dis15* mutant, both SL signalling genes and *AtNAP* respond rapidly to rac-GR24 treatment, indicating these genes were SL-inducible.

**Fig. 9.**
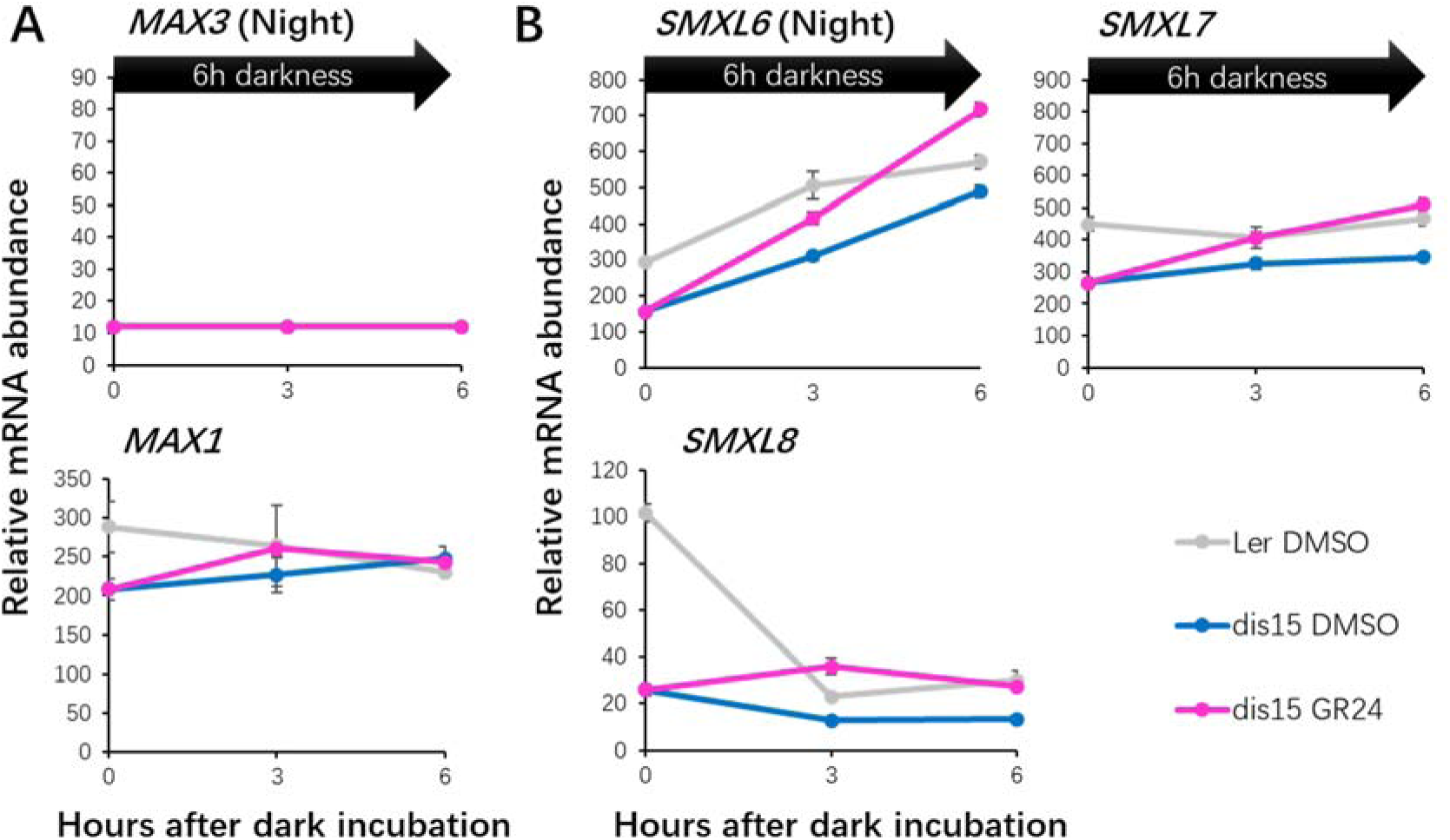
Transcript abundance changes of strigolactone pathway genes during the regular night. (A) Strigolactone (SL) biosynthetic genes; (B) SL signalling genes. Transcript abundance of each gene was quantified using nCounter analysis on RNA isolated from detached WT or *max1-5*/*dis15* inflorescences (n = 3 samples, > 4 inflorescences from independent plants per sample) that were treated with 1% DMSO for WT and 1% DMSO/rac-GR24 for *max1-5*/*dis15* and incubated in the dark for 0 h, 3 h and 6 h. Transcript abundance was normalised to geometric mean of *PP2AA3*, *ACT2* and *MON1*. Data are mean ± standard error. Statistical significance of gene expression for all samples is listed in Supplementary Table S2. Figure legends apply to all figures.

### GR24 suppresses the transcript abundance of SnRK1-related genes in *max1-5*/*dis15* during an extended night

We next determined the effect of extended darkness (i.e. darkness that surpassed the anticipated night period) on carbon status markers, senescence-markers, and SL biosynthesis and signalling genes. In WT, at 4 h of extended night (i.e. 12 h of dark treatment), transcript abundance of *AKINβ1* and *bZIP63* was substantially increased (Fig. 10A; Supplementary Table S2). This was consistent with the WT inflorescences experiencing carbon starvation as has been reported for rosette leaves exposed to a 4 h extended night (Usadel *et al*., 2008). Transcript abundance of *ANAC092* (Kim *et al*., 2009) and *AtNAP* (Guo and Gan, 2006) were also significantly increased at this time (Fig. 10B). However, transcript abundance of *MAX1* was not increased by the 4 h night extension and *MAX3* abundance remained undetectable, suggesting SL biosynthesis was not occurring. By 10 h of extended night (18 h of dark treatment), *MAX1* transcript abundance had still not changed, but that of *MAX3* had increased suggesting SL biosynthesis had started (Fig. 10C). The three *SMXL* genes were differentially expressed during the extended night (Fig. 10D). *SMXL8* transcript counts were undetectable at both 4 h and 10 h of extended night; *SMXL6* transcript abundance was increased at 4 h but then declined; and *SMXL7* started to increase at 10 h of extended night in a pattern strikingly similar to *MAX3*.

**Fig. 10.**
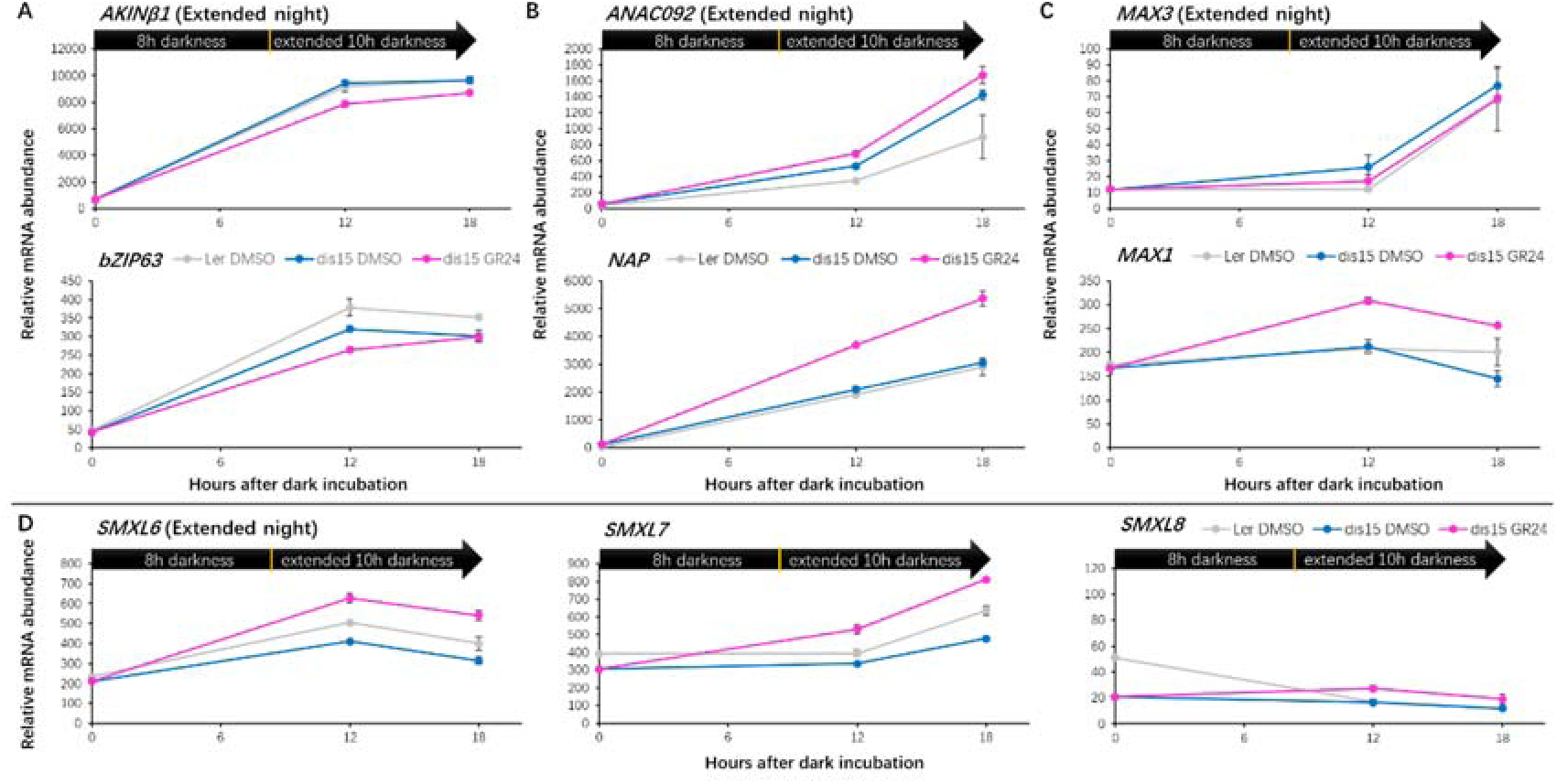
Transcript abundance changes of sugar, senescence and strigolactone pathway genes during the extended night. (A) SnRK1-related genes; (B) functional senescence regulators; (C) strigolactone (SL) biosynthetic genes; (D) SL signalling genes. Transcript abundance of each gene was quantified using nCounter technology on RNA isolated from detached WT or *max1-5*/*dis15* inflorescences (n = 3 samples, > 4 inflorescences from independent plants per sample) that were treated with 1% DMSO for WT and 1% DMSO/rac-GR24 for *max1-5*/*dis15* and incubated in the dark for 0 h, 12 h and 18 h. Transcript abundance was normalised to geometric mean of *PP2AA3*, *ACT2* and *MON1*. Data are mean ± standard error. Statistical significance of gene expression for all samples is listed in Supplementary Table S2.

We then determined how the patterns of expression of the above genes were affected by SL deficiency by examining their transcript accumulation in the *max1-5*/*dis15* mutant. Overall the patterns of accumulation of carbon-status-related, senescence-marker genes and SL biosynthesis and signalling genes in the mutant were very similar to what was seen in WT over the 10 h extended night (Fig. 10A, B, C), suggesting that their pattern of regulation was not controlled by SL. Interestingly, when the mutant was treated with rac-GR24, the transcript abundance of sugar-related genes *AKINβ1* and *bZIP63* were suppressed significantly by rac-GR24 at 4 h and not at 10 h of extended night. By contrast, the transcript abundance of the two senescence-related genes *ANAC092* and *AtNAP* was elevated at 12 h by rac-GR24 and so was *MAX1*. All three *SMXL* genes were upregulated by rac-GR24 at both time points (Fig. 10D) as observed during the regular night.

Taken together, the nCounter profiling study has highlighted a temporal sequence of events whereby markers of carbon deprivation and senescence regulation first increase followed within hours by markers for SL production. Secondly, GR24 treatment of the *max1-5*/*dis15* mutant indicated that SL acts to promote transcription of senescence controlling genes and suppress transcription of SnRK1-related genes.

## Discussion

### SLs regulate sepal senescence in *Arabidopsis*

SLs are plant hormones that affect seed germination (Toh *et al*., 2012), shoot branching (Gomez-Roldan *et al*., 2008; Umehara *et al*., 2008), root architecture (Kapulnik *et al*., 2011; Ruyter-Spira *et al*., 2011; Rasmussen *et al*., 2012) Here we demonstrate that SLs regulate floral organ senescence. We identified two mutants, *max1-5*/*dis15* (MAX1-G469R) and *d14-6*/*dis9* (D14-S97F), which have novel loss-of-function alleles in the SL biosynthetic gene *MAX1* and receptor *AtD14*, respectively. Both mutants exhibited delayed sepal senescence *in planta* and in detached inflorescences upon dark or long day treatment. Such phenotypes were rescued in the *max1-5*/*dis15* but not *d14-6*/*dis9* mutant by treatment of the detached inflorescences with rac-GR24. Transcript abundance of senescence marker genes was supressed in the two mutants compared with in WT in the dark. These results indicate that SLs regulate both developmental and dark-induced sepal senescence.

### Gly-469 of MAX1 is essential for catalysing the conversion of CL into CLA

SLs are synthesised from β-carotene (Alder *et al*., 2012). SL biosynthesis starts with conversion of all-trans-β-carotene into CL, a common precursor of all SLs (Seto *et al*., 2014; Wang and Bouwmeester, 2018). In *Arabidopsis*, this requires the sequential activities of the carotenoid isomerase DWARF27 (D27) and two carotenoid cleavage dioxygenases CCD7/MAX3 and CCD8/MAX4 (Alder *et al*., 2012). Following this, MAX1 oxidises CL into CLA (Abe *et al*., 2014), which is further converted to other SL-like compounds by other downstream enzymes, such as LATERAL BRANCHING OXIDOREDUCTASE (LBO) (Brewer *et al*., 2016).

*MAX1* encodes a CYP711A1 protein that belongs to the cytochrome P450 superfamily (Booker *et al*., 2005). The MAX1 G469R substitution we identified occurs at the last residue in the highly conserved cysteine haem-iron ligand signature of the cytochrome P450 superfamily, which is just two amino acids C-terminal to the absolutely conserved Cys at position 467 of MAX1. To date, no crystal structure of MAX1 has been reported. However, a 3D model based on the closest homologous structure, the human microsomal P450 CYP3A4, revealed that Gly-469 packs against the haem cofactor in the binding pocket and is close to the haem-iron ligand Cys-467 (Fig. 6). Since Gly-469 is very close to this Cys, substitution of the smallest amino acid Gly to the larger Arg likely introduces steric clashes within the haem binding pocket and loss of activity of *max1-5*/*dis15* (MAX1-G469R). Interestingly, the G469 residue is invariant in MAX1 orthologues and its closely related proteins in Metazoa, Bacteria and Archaea [Fig. 6A; Supplementary Fig. S8; (Challis *et al*., 2013)], whereas in the wider cytochrome P450 family in rare instances this residue is replaced by Ala, which suggests that this small nonpolar amino acid may not introduce steric clashes in the wider family of cytochrome P450 proteins. This is consistent with the equivalent Gly to Ala substitution not affecting activity of the *Arabidopsis* cytochrome P450 CYP83B1, a modulator of auxin homeostasis (Barlier *et al*., 2000; Bak *et al*., 2001). Our transient expression assay in *N. benthamiana*, and successful genetic complementation of the *max1-5*/*dis15* mutant with MAX1-G469A demonstrated that the Ala substitution did not inactivate MAX1 function, whereas substitution with arginine did. The high content of CL (the substrate of MAX1) and strongly reduced production of CLA hexose conjugates in the leaves infiltrated with the *MAX1*-G469R construct are consistent with accumulation of CL previously observed for the *Arabidopsis* T-DNA insertion mutant *max1-4* (Seto *et al*., 2014) and indicates that conversion of the MAX1 substrate is prevented. Thus, we conclude that G469 is an important amino acid for MAX1 function, though it can be replaced by Ala.

### Substitution of catalytic Ser-97 affects activity of AtD14

The *d14-6*/*dis9* mutant displayed delayed dark-induced senescence and bushy phenotypes. SL perception and signalling require the D14 receptor. D14 belongs to the protein superfamily of α/β-fold hydrolases (Arite *et al*., 2009). The crystal structures of *At*D14 and its orthologs in petunia (*Ph*DAD2) and rice (*Os*D14) reveal that they have a hydrophobic substrate-binding pocket containing a Ser-His-Asp catalytic triad essential for hydrolase activity (Hamiaux *et al*., 2012; Kagiyama *et al*., 2013; Zhao *et al*., 2013). Unlike the classical hormone receptors that non-covalently and reversibly bind to hormone molecules, the crystal structure of the SL-induced AtD14–D3–ASK1 complex reveals that AtD14 binds to SL and hydrolyses it into a CLIM (Yao *et al*., 2016). It is under debate whether this hydrolysis is required for signalling to occur (Seto *et al*., 2019). Experiments with the F-box protein D3, a rice ortholog of *Arabidopsis* MAX2 showed that SL triggers SL signalling by enabling AtD14 to bind to MAX2 to recruit repressors (e.g. *Arabidopsis* SMXL6/7/8) for degradation through the 26S proteasome (Jiang *et al*., 2013; Zhou *et al*., 2013; Wang *et al*., 2015). In the *d14-6*/*dis9* mutant, Ser-97 in the catalytic triad was replaced by Phe. This produced phenotypes similar to the null mutant *d14-1* suggesting loss of activity of D14 (Waters *et al*., 2012; Ueda and Kusaba, 2015). This is consistent with mutation of this residue (atd14:S97A) affecting binding of the receptor to the SL analogue GR24 and abolishing hydrolase activity (Abe *et al*., 2014). Thus, it is probable that the S97F mutation we identified causes complete loss of D14 activity.

### SLs regulate dark-induced inflorescence senescence in association with a change of carbon status during the extended night

SLs affect various stress responses including drought, high salinity, nutrient deficiency and light-deprivation/darkness (Bu *et al*., 2014; Ha *et al*., 2014; Yamada *et al*., 2014; Ueda and Kusaba, 2015; Yamada and Umehara, 2015). Our findings suggest that SLs also regulate senescence of *Arabidopsis* sepals under energy-deprivation conditions. We previously demonstrated that the energy-deprivation resulting from prolonged darkness causes a reduction in soluble sugars and transcriptional changes of sugar-responsive genes in detached inflorescences (Trivellini *et al*., 2012). This suggests an interaction between SLs and sugar signalling in controlling dark-induced inflorescence senescence. Crosstalk between SLs and sugar signalling has been reported for regulation of shoot branching and seedling establishment in *Arabidopsis* (Li *et al*., 2016; Otori *et al*., 2017). To investigate the interaction between SLs and sugar signalling in controlling inflorescence senescence, we harvested WT and SL deficient *max1-5*/*dis15* inflorescences at the start of the dark cycle and examined the transcriptional changes of selected genes in SL, sugar, and senescence pathways after different times of dark incubation.

During the night, SnRK1s have a crucial role in the sugar-dependent regulation of transcriptional response (Blasing *et al*., 2005; Baena-Gonzalez *et al*., 2007; Usadel *et al*., 2008). It is therefore not surprising that in WT inflorescences SnRK1-related genes *AKINβ1* (a subunit of SnRK1) (Li *et al*., 2009) and *bZIP63* (one of the direct targets of SnRK1) (Mair *et al*., 2015) were both up-regulated at 3 h of dark incubation. Such early upregulation was also observed in *Arabidopsis* rosettes and attributed to starch decline during the normal night (Blasing *et al*., 2005). Interestingly, in WT inflorescences key senescence regulators *ANAC092* (Kim *et al*., 2009) and *NAP* (Guo and Gan, 2006) were also up-regulated at 3 h. However, in rosette leaves the decline in sugar content during the night does not cause starvation (Blasing *et al*., 2005; Usadel *et al*., 2008) and therefore it is unlikely that up-regulation of these marker genes in the inflorescences was indicative of senescence initiation. Moreover, since expression of both genes also depends on sugar content and circadian clock (Blasing *et al*., 2005; Usadel *et al*., 2008; Kim *et al*., 2018a; Song *et al*., 2018), their transcriptional changes more likely result from one or both of these cues. Because of technological limitations, precise quantification of SL content in *Arabidopsis* is extremely difficult (Seto *et al*., 2014; Lv *et al*., 2018). Thus, we used the transcriptional changes of selected SL biosynthetic genes to indirectly estimate SL content (Li *et al*., 2018). We found in WT during the normal night that *MAX1* abundance was unchanged and *MAX3* counts were below the set threshold for detection. This suggests that SLs are not synthesised during the regular night. We also examined transcriptional changes of SL signalling genes *SMXL6*, *SMXL7* and *SMXL8* to help determine timing of SL response. These three *SMXL*s are functionally redundant in controlling *Arabidopsis* shoot branching (Soundappan *et al*., 2015; Wang *et al*., 2015). We considered their involvement in senescence regulation because senescence and branching phenotypes were tightly linked in*max1-5*/*dis15* and *d14-6*/*dis9* mutants. *SMXL7* transcript abundance was unchanged in WT during the normal night, consistent with expression patterns of the biosynthetic genes. However, *SMXL6* transcript increased significantly at 3 h and *SMXL8* transcript declined rapidly to become undetectable at 3 h and onwards. This raises the possibility that distinct cues affect transcription of these three *SMXL*s. Transcript abundance of the three *SMXLs* was lower in *max1-5*/*dis15* inflorescences but was restored by rac-GR24 treatment. These findings suggest that the regular night causes sugar levels to decline without affecting SL biosynthesis.

The extended night commences when the regular night ends and it was previously shown that in *Arabidopsis* rosettes, carbon became severely limited 4 h into the extended night (Usadel *et al*., 2008). *SnRK1*s, have key roles in regulating transcriptional reprogramming during energy-deprivation (Baena-Gonzalez *et al*., 2007; Usadel *et al*., 2008). Consistent with these observations, we found that transcript abundance of *AKINβ1* and *bZIP63* increased more in WT at 12 h (4 h into extended night) than during the regular night. The sugar-responsive and senescence-controlling genes *ANAC092* and *NAP* also showed further increased transcript abundance 12 h. Then at 18 h (10 h into extended night), *AKINβ1* and *bZIP63* did not show further transcriptional changes. This could be because sugar depletion in the inflorescence is alleviated by metabolic readjustment, similar to what has been found in rosette leaves, where sugar (Glc-6-P and Fru-6-P) levels were partially recovered from 4 h onward in the extended night (Usadel *et al*., 2008). By contrast, *ANAC092* and *NAP* transcript abundance continued to rise. This raises the possibility that at 18 h the *ANAC092* and *NAP* gene products started to initiate senescence to prevent critically low sugar levels. In the *max1-5*/*dis15* mutant, rac-GR24 treatment transiently repressed *AKINβ1* and *bZIP63* at 12 h, while *ANAC092* and *NAP* were up-regulated. Here, the SL/GR24 treatment may induce senescence – and associated sugar recovery - earlier, alleviating the need for metabolic readjustment. Alternatively, the SL/GR24 may transcriptionally repress key energy-deprivation regulators directly, which then prevents recovery of sugar shortage and as a necessity promotes senescence as another nutrient recovery process. Regardless, these findings are consistent with the idea that SL promotes senescence to restore sugar shortage. SLs may not play a major role in metabolic adjustment until 18 h, when the *MAX3* and *SMXL7* transcripts were up-regulated in the WT. *SMXL7* has been found to exhibit higher transcript abundance than *SMXL6* and *SMXL8* in senescent leaves (Stanga *et al*., 2013). Thus, the up-regulation of *SMXL7* at 18 h may indicate a senescence regulatory role for this gene. However, we could not ignore the possibility that *SMXL6* is also involved in regulating senescence because of its significantly changed transcript abundance at 12 h and 18 h as well as its important role in regulation of shoot branching (Soundappan *et al*., 2015; Wang *et al*., 2015).

We previously found during 24-h of dark treatment that soluble sugar and Chl contents reduced substantially in the inflorescence (Trivellini *et al*., 2012). The dark treatment was accompanied with transcriptome changes of carbon-deprivation and senescence-related genes including *AKINβ1*, *bZIP63*, *ANAC092* and *NAP* (Trivellini *et al*., 2012). In this study, expression of senescence-specific marker gene *SAG12* started to increase at 48 h in WT suggesting that senescence was well underway at that time. The transcript abundances of *SAG12*, *ANAC092* and *NAP* increased more strikingly at 72 h and similar increases were found for the SL biosynthetic genes *MAX3*, *MAX4* and *MAX1*. In the SL mutants, the senescence process was slowed down and expression of *ANAC092* and *SAG12* lower than in the WT (Fig. 3). This demonstrates that SLs contribute to the senescence of dark-detached inflorescences and is consistent with what has been observed during leaf senescence (Ueda and Kusaba, 2015). The expression pattern of signalling gene *SMXL7* showed the strongest correlation with that of SL biosynthetic genes and senescence marker genes, which suggests involvement in senescence regulation. However, the decreased transcript abundance of *SMXL6* after 24 h suggests it may play a more important role in early stages of senescence.

Overall, our results suggest an intricate relationship among sugar starvation, senescence and SL biosynthesis and signalling in excised inflorescences. This is supported by a recent study finding that the SL-induced senescence of bamboo leaves was supressed by exogenous sugar treatment (Tian *et al*., 2018). SLs do not appear to have a major role in the inflorescences during the normal night but are synthesised during the extended night, perhaps in response to sustained low sugar content and consequent senescence initiation. It seems thus that SLs play an important role in the regulation of nutrient redistribution from leaves and leaf-like organs to other organs of the plant by promoting senescence progression.

## Materials and methods

### Plant growth conditions and mutant analysis

EMS-mutated seeds (M2) from *Arabidopsis thaliana* ecotype Landsberg *erecta* (L*er*-0) background were purchased from LEHLE Seed Company (Round Rock, TX, USA; www.lehleseeds.com). Seeds of *Arabidopsis* ecotype Columbia (Col-0), L*er*-0 and EMS mutants were germinated and grown in a temperature-controlled growth chamber set at 21°C with 65% relative humidity and under a 16 h/8 h light (200 μM photons m^−2^s^−1^; Gro-Lux and cool-white fluorescent lamps)/dark cycle (long day) unless otherwise stated. For the *in planta* assay, plants were grown in a temperature-controlled growth cabinet (Contherm Model CAT 630, Wellington, New Zealand), 20-22°C with 60% relative humidity at a light intensity of ∼180 μE with Metal Halide lamps under long day conditions. Plants were screened by using *Arabidopsis* inflorescence degreening assay as described in Hunter *et al*. (2018). For long day treatment, similar procedures were used with the exception that inflorescences were placed in a transparent container that was covered with transparent film and incubated under long day. For SL treatment, inflorescences were treated with rac-GR24 (Chiralix, Nijmegen, The Netherlands), which was dissolved in pure DMSO and diluted to different concentrations for experiments, or 1% DMSO (mock). Mutant selection and genetic analysis are described in Jibran *et al*. (2015). Selected mutants were backcrossed twice to the WT L*er*-0 prior to analysis and genetic complementation.

### Chlorophyll analysis

Chl was extracted from single inflorescences using the protocol described in Jibran *et al*. (2015) with the following changes: fresh samples were used for extraction and the samples were incubated in the dark at 4°C for 4 days after adding ethanol. The Chl retention at day 3 was expressed as percentage of day 0.

### HRM-based mapping

High Resolution Melting (HRM)-analysis was performed using the genomic DNA isolated from the leaves of F2 plants that displayed the *dis* phenotype. Genomic DNA for HRM-analysis was isolated by SlipStream Automation (www.slipstream-automation.co.nz) or according to the protocol of Dellaporta *et al*. (1983). The causal mutations were coarse mapped to chromosome 3 using primers 3-1 to 3-4 for *dis9* and chromosome 2 using primers 2-2 to 2-6 for *dis15* (Hunter *et al*., 2018). The additional primers used for fine mapping of *dis9* were designed using Primer3web version 4.1.0 (Koressaar and Remm, 2007; Untergasser *et al*., 2012) according to the settings described in Jibran *et al*. (2015) and are listed in Supplementary Table S3. HRM reaction was prepared using HRM master mix (Roche Diagnostics, Mannheim, Germany) and the PCR was carried out with LightCycler 480 instrument (Roche Diagnostics) following the procedures described in Jibran *et al*. (2015) and Hunter *et al*. (2018).

### Genomic DNA isolation and whole genome sequencing analysis

Genomic DNA for WGS was isolated from *dis9* using the Plant Genomic DNA Mini Kit (Geneaid, Taiwan, China) and from *dis15* using the method of Lutz *et al*. (2011). Genomic DNA at a concentration of 100 ng/μL in 1× TE buffer was sent to Macrogen (http://dna.macrogen.com) for 100 bp paired end sequencing on an Illumina HiSeq2000 sequencer. The trimmed reads were aligned to the L*er*-0 reference using Geneious desktop software (Kearse *et al*., 2012) or Bowtie 2 software (Langmead and Salzberg, 2012) for SNP detection between the mutant and WT.

### *MAX1* and *AtD14* constructs for complementation

For *MAX1* constructs, the fragment of *MAX1*-WT (consisting of the 1293 bp region upstream of the start codon, the 2420 bp genomic sequence, and a 185 bp region downstream of the stop codon) or *MAX1-*G469A (with a G to C mutation at position 1406 relative to the start codon of the *MAX1* coding sequence causing a G469A substitution of MAX1 protein) was amplified from genomic DNA with indicated primers (Supplementary Table S4) and was then cloned into the binary vector *pGreen 0229* (Hellens *et al*., 2000) via restriction digestion with *EcoR*I and *Not*I enzymes (Roche Diagnostics, Mannheim, Germany). The *AtD14* construct (comprising 553 bp sequence upstream of the start codon and 804 bp of the coding sequence) was provided by Pilar Cubas (Centro Nacional de Biotecnología/Consejo Superior de Investigaciones Científicas; (Chevalier *et al*., 2014)). The positions of the mutations and the cloned regions were assigned based on the Col-0 gene and protein reference sequences from TAIR (www.arabidopsis.org). These constructs were transformed into *A. tumefaciens* GV3101 by electroporation and were then transformed by agroinfiltration to homozygous mutant plants using the floral dip method (Zhang *et al*., 2006).

### RNA isolation and qRT-PCR analysis

Total RNA was isolated with the Quick-RNA™ MiniPrep kit (Zymo Research, Irvine, CA, USA) according to the instruction manual (with on-column DNase treatment). cDNA was synthesised from RNA (500ng) using iScript™ Reverse Transcription Supermix for RT-qPCR (BIO-RAD, Hercules, CA, USA). The cDNA was diluted 10-times for quantitative PCR (qRT-PCR) analysis. qRT-PCR reaction was prepared using a LightCycler® 480 SYBR Green I Master kit (Roche Diagnostics, Mannheim, Germany) and the PCR was performed with a LightCycler® 480 Instrument II (384-well; Roche Diagnostics) according to the manufacturer’s instructions on three biological replicates (each with 4 to 6 pooled inflorescences from individual plants). Four technical replicates were performed for each biological replicate. Primers were designed using QuantPrime online software (Arvidsson *et al*., 2008) and are listed in Supplementary Table S5. The Cp value was calculated using the algorithm of “Abs Quant / 2nd Derivative Max” present in LightCycler® 480 Software (version 1.5). Data were normalized to the reference gene *PP2AA3* (*At1g13320*) (Czechowski *et al*., 2005) and relative transcriptional changes were calculated using the ΔΔCt method (Livak and Schmittgen, 2001; Dvinge and Bertone, 2009).

### nCounter analysis

Transcriptional analysis was performed using the nCounter Analysis System (NanoString, Seattle, WA, USA) (Geiss *et al*., 2008). Two sets of gene-specific probes (along with a reporter probe and a capture probe) were designed by NanoString Support and their sequences are listed in Supplementary Table S6. Total RNA (300 ng per sample) was hybridised using nCounter PlexSet-24 Reagent Pack according to the “PlexSet™ Reagents User Manual”. After hybridisation, samples were vertically pooled and were placed on the automated nCounter Prep Station (NanoString, USA) for purification and were immobilised in the cartridge. This cartridge was then transferred to the nCounter Digital Analyzer for data collection. Data analysis was performed with nSolver™ 4.0 Analysis Software according to user manual. All samples passed the quality control. The background thresholding was set to “12” according to the count value of the internal negative control. Positive control normalization was carried out by using the geometric mean of the Top 3 positive counts. Reference gene normalization was calculated using the geometric mean of counts for the three reference genes *PP2AA3*/*At1g13320*, *ACT2*/*At3g18780* and *MON1*/*At2g28390* (Czechowski *et al*., 2005).

### *In silico* analysis

Sequence alignments were performed using Geneious desktop software (Kearse *et al*., 2012). The three-dimensional structure of AtD14 was obtained from the SL-induced AtD14-D3-ASK1-complex (PDB: 5HZG) (Yao *et al*., 2016). The homology model of MAX1 was calculated using I-TASSER Online Server (https://zhanglab.ccmb.med.umich.edu/I-TASSER/) (Zhang, 2008; Roy *et al*., 2010; Yang *et al*., 2015). The 3D images were prepared with CCP4MG (McNicholas *et al*., 2011).

### Plasmid construction for transient expression assay

The *Arabidopsis* Col-0 based SL biosynthetic genes (*D27*, *MAX3*, *MAX4* and *MAX1*) were cloned as described in Zhang *et al*. (2014). The L*er*-0 based *MAX1* gene (*MAX1*-WT, *MAX1*-G469R or *MAX1*-G469A) was cloned using the same protocol but with the primers listed in Supplementary Table S7.

### Transient expression in leaves of *Nicotiana benthamiana*

To characterise the enzymatic activity of MAX1 and the genetic variants described herein we used transient expression in *N. benthamiana* essentially as described in Zhang *et al*. (2014) except that *A. tumefaciens* were resuspended in 50 mM MES (2-Morpholinoethanesulfonic acid hydrate) (Duchefa, Haarlem, The Netherlands) - KOH buffer (pH 5.6) containing 2mM NaH_2_PO_4_ (Merck, Darmstadt, Germany), 100 µM acetosyringone (Sigma-Aldrich, St. Louis, MO, USA) and 0.5% glucose (MP Biomedicals, France) to a final OD_600_ of 0.5. Instead of *OsD27*, *OsCCD7* and *OsCCD8* that were shown by Zhang *et al*. (2014) to result in the production of CL upon transient expression in *N. benthamiana* we used *AtD27*, *MAX3* and *MAX4*. By co-infiltrating *MAX1* and genetic variants with these genes we can study the conversion of CL to CLA. Infiltration was performed using 4-week-old *N. benthamiana* plants. For each gene combination, six individual plants were used as biological replicates.

### Analysis of carlactone and carlactonoic acid in *Nicotiana benthamiana*

Samples were analysed for the presence of CL and CLA (using UPLC-LC-MS/MS) and CLA conjugates (untargeted analysis using UHPLC-qTOF-MS). For both analyses, 200 mg fine-ground *N. benthamiana* leaves were extracted in 2 mL of ethyl acetate, using GR24 (5 pmol) as internal standard. Samples were vortexed and centrifuged for 20 min at 2000 rpm. at 4°C. The supernatant was dried *in vacuo*. Prior to mass analysis samples were reconstituted in 100 µl of 25% acetonitrile/water (v/v) and filtered using a micro-spin 0.2 µm nylon membrane filter (Thermo Fisher Scientific, Waltham, MA, USA).

The targeted analysis of CL and CLA was performed by Acquity UPLC system (Waters, Milford, MA, USA) coupled to a Xevo® TQ-XS triple-quadrupole mass spectrometer (Waters MS Technologies, Manchester, UK) with electrospray (ESI) interface. Samples were injected onto a reverse-phase UPLC® column Acquity BEH C18 (2.1×100mm, 1.7 µm, Waters), thermostated at 45°C. The retention of analytes was controlled by gradient elution of 15 mM formic acid in water (A) and 15 mM formic acid in acetonitrile (ACN, B) at a flow rate 0.4 ml/min. The 10 min linear gradient started by isocratic elution 0-0.5 min with 5% B, increased to 60% B in 1.5 min and to 90% B in next 5.3 min. The column was washed for 1.5 min with 90% B and equilibrated for initial conditions for 1.5 min. The eluate was introduced in the ESI ion source of the triple quadrupole MS analyser, operating in both positive and negative mode at following conditions: capillary voltage (1.2 kV), ion source/desolvation temperature (150/600°C), desolvation/cone gas flow (1000/150 L.h^−1^), cone voltage (20-25 V) and collision energy (18-25 eV). MS data were recorded in multiple reaction monitoring mode (MRM) of four characteristic transitions for each of compounds. MassLynxTM software package (version 4.2, Waters, Milford, MA, USA) was used to operate the instrument, and acquire and process MS data.

### Detection and quantification of carlactonoic acid conjugates by UHPLC-qTOF-MS

The above described *N. benthamiana* leaf extracts were also analysed on ultra-high-performance-liquid chromatography-quadrupole-time-of-flight-mass-spectrometry (UHPLC-qTOF-MS) consisting of an Agilent 1290 LC coupled to a Bruker Daltonics microTOF-Q mass spectrometer (Bremen, Germany). The LC was equipped with a KINETEX® XB-C18 column (2.1 mm x 100 mm, 2.6 μm; Phenomenex). Mobile phase A consisted of 5% ACN in water and 0.1 % formic acid, whereas mobile phase B consisted of 95% ACN and 0.1% formic acid. The gradient was 0-3 min (isocratic at 95% A), 3-35 min (increase to 100% B), 40-41 min (decrease to 95% A) and column equilibration for 9 min at initial conditions (95% A). The chromatographic run lasted 40 min with a flow rate of 0.2 mL min^−1^. The mass spectrometer was operated in negative mode. The mass spectrometer’s settings were: dry gas flow rate 8 L min^−1^ at 220°C, capillary voltage 3,8 kV, collision energy 10 eV and collision RF 1200 Vpp. The QTOF was operated with the *m/z* range set from 50 to 1500 Da. The injection volume was 10 μL. Acquisition of LC-MS data was performed using Bruker DataAnalysis 4.3.

### Statistical analysis

Statistical analysis was performed with GenStat 17^th^ Edition (a VSNI product: https://www.vsni.co.uk/software/genstat/). One-way ANOVA (Fisher’s protected LSD test *p*<0.05) was used to determine the statistical significances for the Chl data, qRT-PCR data and nCounter data for a period of 72-h of dark treatment in WT Ler-0. A linear mixed model was used to determine the differences for the data of a period of 6-h and 18-h treatments in both WT and *max1-5*/*dis15* (with rac-GR24 or 1% DMSO treatment). Comparisons among means were made using least significant differences at *p*=0.05 (5% LSD). CL and CLA data were analysed using one-way ANOVA. To equalise the variances, the variables were log-transformed prior to analysis. Comparisons among means were made using 5% Fisher’s LSDs (least significant differences).

### Accession numbers

Gene and protein accession numbers are listed in Supplemental Table S8.

## Supplemental Data

**Supplemental Fig. S1:** The dwarf and bushy phenotypes of *dis9* and *dis15 in planta*.

**Supplemental Fig. S2:** Delayed sepal degreening of *dis9* and *dis15 in planta* and in detached inflorescences held in the long day conditions (16 h/8 h light/dark cycle)

**Supplemental Fig. S3:** Mapping and cloning of the *dis9* and *dis15* locus.

**Supplemental Fig. S4:** Degreening of inflorescences treated with 1% DMSO (mock) and rac-GR24 at 1, 5, 25, 50 µM.

**Supplemental Fig. S5:** Sepal degreening of inflorescences treated with 1% DMSO (mock) and 5 µM of rac-GR24.

**Supplemental Fig. S6:** Genetic complementation of *dis9*.

**Supplemental Fig. S7:** Complementation of *dis15* with *MAX1* WT and *MAX1*-G469A.

**Supplemental Fig. S8:** Sequence alignment of *Arabidopsis* MAX1 with its putative orthologs, related proteins and other cytochrome P450 proteins with the haem-iron ligand signature.

**Supplemental Fig. S9:** Identification of CLA conjugates in *N. benthamiana* leaves infiltrated with different combinations of SL biosynthetic genes.

**Supplemental Fig. S10:** Transcript abundance of *MAX1* in leaves of *N. benthamiana* infiltrated with constructs harboring *MAX1*-WT, -G469R or -G469A.

**Supplemental Fig. S11:** Transcript abundance changes of SL pathway and senescence-related genes

**Supplemental Fig. S12:** Transcript abundance changes of sugar- and senescence-related genes during the regular night

**Supplemental Fig. S13:** Transcript abundance changes of sugar- and senescence-related genes

**Supplemental Table S1:** Pearson’s chi-squared test for F2 progenies of *dis9* and *dis15*

**Supplemental Table S2:** Statistical analysis of nCounter data for 0, 3, 6 h and 0, 12, 18 h treatments

**Supplemental Table S3:** HRM primers for fine mapping of *dis9*

**Supplemental Table S4:** Primers used for cloning MAX1 and MAX1-G469A for genetic complementation

**Supplemental Table S5:** Primers for qRT-PCR analysis

**Supplemental Table S6:** Probes for nCounter analysis

**Supplemental Table S7:** Primers used for MAX1-related constructions for Agro-infiltration in *N.benthamiana*

**Supplemental Table S8:** Gene and protein accession numbers

## Acknowledgements

We acknowledge the Joint Graduate School of Horticulture and Food Enterprise for a doctoral scholarship to XX and Plant & Food Research Strategic Science Investment fund: 1972 – “Breeding Technology Development” for financial assistance to conduct this research. YW was supported by the Chinese Scholarship Council, LD by the EU (Marie Curie grant NemHatch, 793795), KF by the Netherlands Organisation for Scientific Research (NWO-ECHO grant 711.018.010) and HB by the European Research Council (ERC Advanced grant CHEMCOMRHIZO, 670211). We thank Ian King for help with growing plants, Kerry Sullivan for gDNA isolation for WGS, David Chagné for mentoring on HRM analysis, Dave Wheeler for help with WGS analysis and Aleksandra Chojnacka for LC-QTOF-MS analysis. We thank Pilar Cubas (Centro Nacional de Biotecnología/Consejo Superior de Investigaciones Científicas) for providing D14 construct for complementation.

